# Control tools to selectively produce purple bacteria for microbial protein in raceway reactors

**DOI:** 10.1101/2020.01.20.912980

**Authors:** Abbas Alloul, Marta Cerruti, Damian Adamczyk, David G. Weissbrodt, Siegfried E. Vlaeminck

**Affiliations:** Research Group of Sustainable Energy, Air and Water Technology, Department of Bioscience Engineering, University of Antwerp, Groenenborgerlaan 171, 2020 Antwerpen, Belgium; Department of Biotechnology, Delft University of Technology, van der Maasweg 9, 2629 HZ Delft, the Netherlands

## Abstract

Purple non-sulfur bacteria (PNSB) show potential for microbial protein production on wastewater as animal feed. They offer good selectivity (i.e. uneven community with high abundance of one species) when grown anaerobically in the light. However, the cost of a closed anaerobic photobioreactor (PBR) is prohibitive for protein production. While open raceway reactors are cheaper, their feasibility to selectively grow PNSB is thus far unexplored. This study developed tools to boost PNSB abundance in the biomass of a raceway reactor fed with volatile fatty acids as carbon source. For oxygen availability as tool, not stirring in the night (i.e. reduced oxygen supply) elevated the PNSB abundance from 8% to 20%. For light availability as tool, a 24-h illumination increased the PNSB abundance from 8% to 31% compared to a 12-h light/12-h dark regime. A reactor run at 2-d sludge retention time at the highest surface-to-volume ratio (10 m^2^ m^-3^ increased light availability) showed productivities up to 0.2 g protein L^-1^ d^-1^ and the highest PNSB abundance (78%). The estimated production cost is €1.9 kg^-1^ dry weight (vs. PBR €11.4 kg^-1^ dry weight). This study pioneered in PNSB-based microbial protein production in raceways, yielding cost efficiency along with high selectivity when avoiding the combined availability of oxygen, COD and darkness.

**Graphical abstract:** 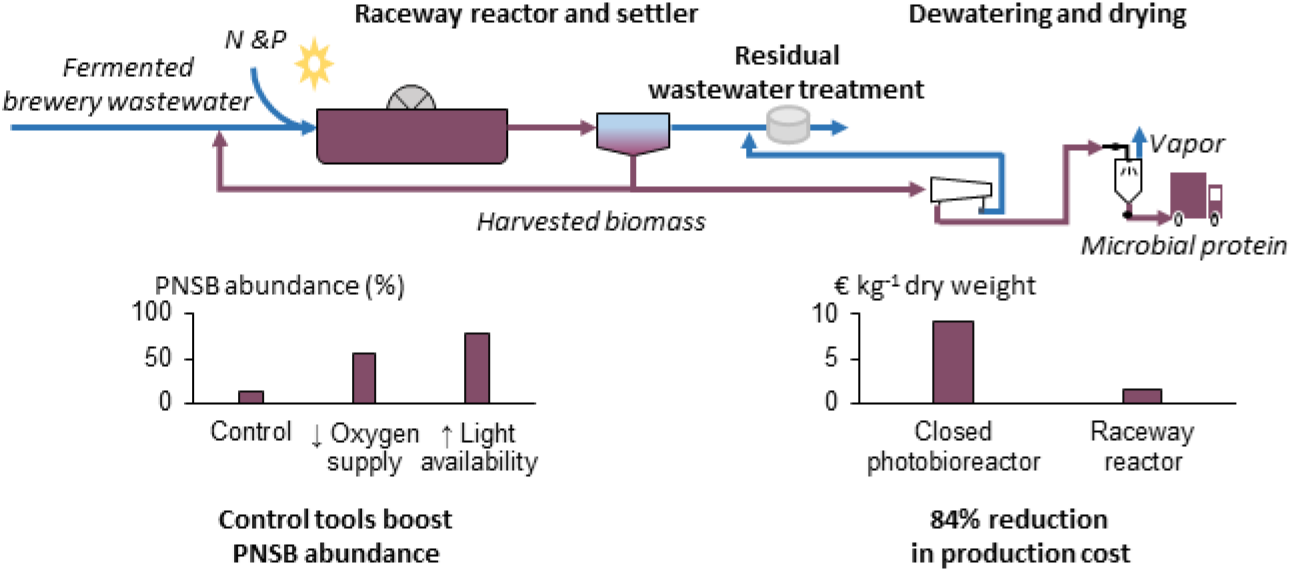

## 1 Introduction

Globally, only 4% of nitrogen and 17% of phosphorus fertilizers applied to the land, are eventually consumed.^1,2^ These inefficiencies in the fertilizer-food chain severely distort the carrying capacity of the Earth, surpassing the planetary boundaries (i.e. safe operating space for sustainability) beyond the zone of uncertainty.^3^ Mitigation can be brought about by upgrading wastewater resources to microbial protein or single-cell protein, which is the use of microorganisms as animal feed ingredient.^4,5^ Resource recovery from food and beverage wastewater is preferred. Brewery wastewater is a key target as it offers a relatively high chemical oxygen demand (COD) concentration (800-9000 mg COD L^-1^) and easiness to prevent fecal contamination.^6^

Upgrading wastewater resources to microbial protein requires either chemo- or photoheterotrophic microorganisms to convert the organic carbon as well as non-axenic production conditions, as it is cost-wise redundant to sterilize vast amounts of water.^4^ Chemoheterotrophs, also known as aerobic heterotrophic bacteria (AHB; i.e. aerobic activated sludge), make use of oxidation reactions for energy generation. These bacteria typically have low biomass yields (0.6 g COD_biomass_ g^-1^ COD_removed_) and high growth rates (2-6 d^-1^).^7^ To date, AHB are pioneering in research, pilot, and full-scale implementations.^8,9^ However, it is challenging to produce an AHB product characterized by an uneven microbial community with a high abundance of one dominant species (i.e. microbial selective production).^8^ Photoheterotrophic cultivation of PNSB may offer such potential, because of their unique ability to grow highly selectively under anaerobic conditions in the light.^10–12^ PNSB are characterized by high biomass yields (0.9-1.1 g COD_biomass_ g^-1^ COD_removed_) and have growth rates between 0.6-3.7 d^-1^.^13–15^ However, compared to AHB, there is a lack of full-scale PNSB facilities for microbial protein production, probably due to costs (expensive photobioreactor; PBR).

To achieve selectivity with PNSB (i.e. uneven microbial community and high abundance of one species), current research has focused on closed PBR such as anaerobic membrane bioreactors,^16^ anaerobic tubular PBR^13,17^ and illuminated anaerobic sequencing batch reactors^18^. These closed PBR only allow the growth of phototrophs and anaerobic chemotrophs. In the case of our previous study on synthetic wastewater, PNSB were able to be selectively produced with an abundance of *Rhodobacter capsulatus* between 93-97% and a low diversity index (exponent Shannon index) between 1.2-1.5 corresponding to an uneven microbial community.^19^ A current cost estimation for PNSB-based protein production on wastewater in a closed PBR amounts to €22 kg^-1^ protein,^13^ which is roughly 11 times higher compared to the price of fishmeal of €2 kg^-1^ protein.^20^.

An economically more interesting case can be made if PNSB were produced in open raceway reactors conventionally used for microalgal processes.^21^ These raceway reactors are open systems with a reactor depth around 20 cm (vs. 6 cm diameter tubular PBR), a surface-to-volume ratio of 5 m^2^ m^-3^ (vs. 22 m^2^ m^-3^ tubular PBR) and are agitated through a paddlewheel (vs. circulation pumps tubular PBR).^21^ Investment costs of a raceway reactor approximate €56 m^-3^ compared to €5,000 m^-3^ for a closed PBR.^22,23^ However, achieving selective PNSB production is more challenging in these reactors, as air continuously enters the system, which enables the proliferation of competing aerobic heterotrophs (i.e. non-PNSB). Moreover, the oxygen concentration in a raceway reactor is zero due to its direct use as electron acceptor, making the growth of anaerobic chemotrophs also possible (e.g. acidogenic microorganisms and sulfate-reducing bacteria; SRB). Currently, there is no published research available on PNSB production with raceway reactors, except for a trial focusing on polyhydroxyalkanoate production.^24^ However, it can be anticipated that the following control tools are essential to maximize PNSB selectivity: (i) limiting the oxygen supply may decrease the growth of aerobic chemotrophs, (ii) increasing the light availability or the illumination period may aide PNSB in their competition for COD with aerobic chemotrophs and anaerobic chemotrophs such as acidogenic microorganisms and SRB, (iii) short sludge retention times (SRT) may washout slower-growing microorganisms such as microalgae, and (iv) limiting the COD-loading rate may decrease the competition with (an)aerobic chemotrophs during the dark period.

The hypothesis of this study was that PNSB can be selectively produced in a raceway reactor provided a good combination of control tools. Batch experiments were first performed to assess the phototrophic and (an)aerobic chemotrophic conversions of PNSB (axenic) and the (an)aerobic chemotrophic conversions of non-PNSB (non-axenic) to understand how they individually would contribute in a raceway reactor. Afterwards, the effect of oxygen supply, light availability, SRT and COD-loading rate was studied on PNSB abundance and community diversity (non-axenic). A raceway reactor was then operated at a SRT of 2 d to understand whether PNSB can be selectively produced over sequential batches and explore the best operational strategy for protein production and COD removal. All experiments were performed using a synthetic medium composed of volatile fatty acids (VFA). The findings of the raceway reactor were finally economically evaluated based on a production cost estimation, and benchmarked with the production of a PNSB biomass in a closed PBR.

## 2 Materials and methods

### 2.1 Inocula and medium

A *Rhodobacter capsulatus* strain, isolated in our previous study,^13^ was used as model PNSB for the axenic flask and non-axenic raceway reactor experiments. This species was selected based on a prior evaluation made between five PNSB cultures, where it showed to have the highest photoheterotrophic growth rate on synthetic wastewater. This species is able to grow photo- and chemoheterotrophically,^10^ which enables examination of different PNSB growth kinetics in raceway reactors. Aerobic activated sludge of a local brewery company (AB InBev, Belgium, Leuven) was used as a proxy for a non-PNSB inoculum.

An adapted VFA-based medium as a proxy for fermented wastewater was used for all experiments, as we argued in a previous study that fermentation prior to protein production will favor the microbial selectivity.^4,13^ The COD concentration was increased to 3 g L^-1^ and contained a defined mixture of acetate, propionate and butyrate on a 1/1/1 COD basis. PNSB grown photoheterotrophically on this VFA mixture have a biomass yield that approximate 1 g COD_biomass_ g^-1^ COD_removed_.^13,16^ This makes it easy to assess the chemoheterotrophic growth of PNSB and of competing non-PNSB, as a lower biomass yield implies oxidation of COD to CO_2_.

### 2.2 Overview of the experiments

Five sets of flask and raceway reactor experiments were performed in this study to explore the conversions of PNSB and the effect of oxygen supply, light availability and SRT on PNSB selectivity (Table 1).

**Table 1.**
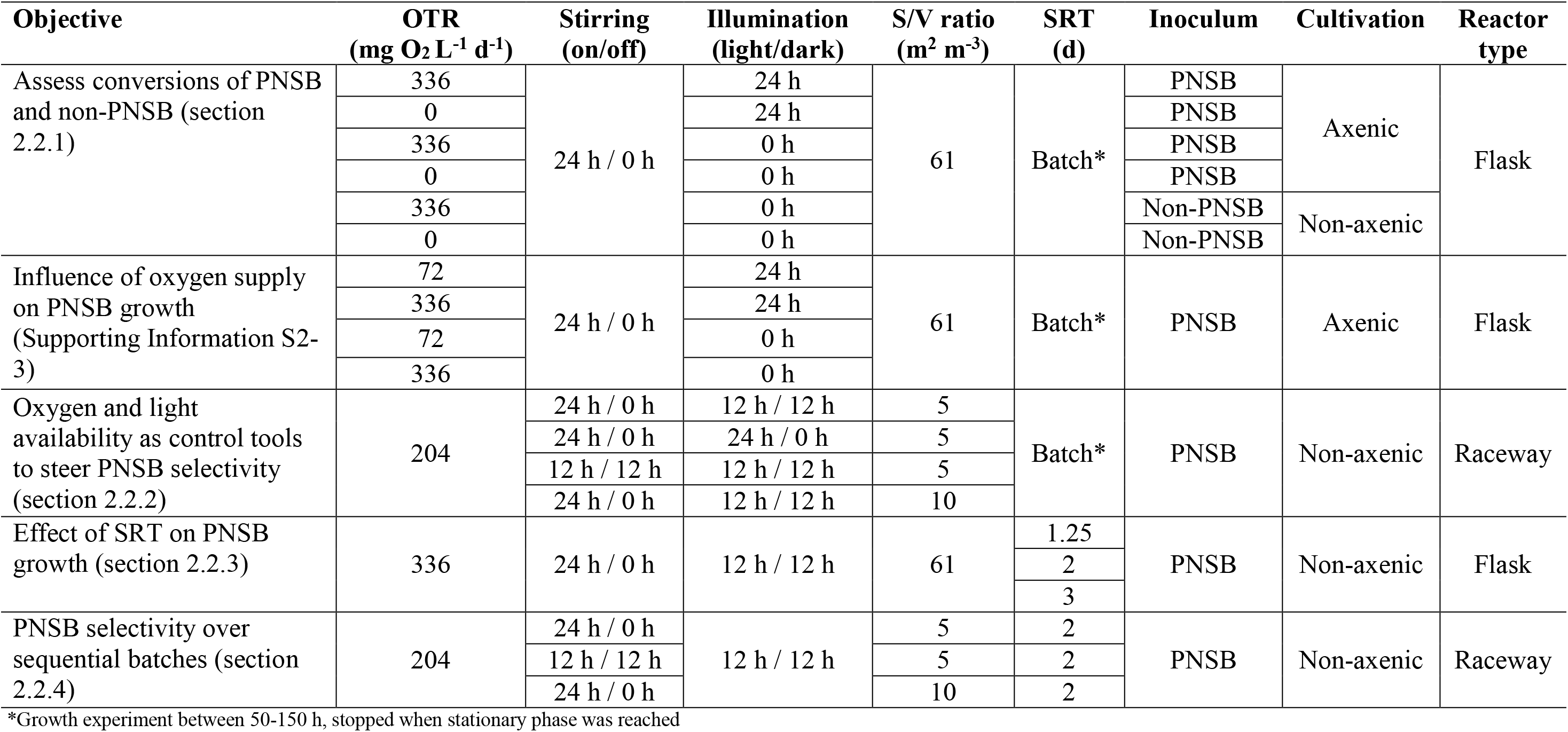
Objectives and experimental setup of five tests to grow a protein-rich PNSB biomass on brewery wastewater. *Rhodobacter capsulatus* was used as purple non-sulfur bacterium (PNSB) inoculum and aerobic brewery sludge as non-PNSB inoculum. Experiments were performed at 28°C. Surface-to-volume (S/V) ratios were calculated based on the illuminated surface area. The flasks were illuminated from the side and the raceway reactor from the top. OTR: oxygen transfer rate; SRT: sludge retention time

#### 2.2.1 Assess the conversions of PNSB and non-PNSB

Flask batch experiments were performed to explore the photoheterotrophic and (an)aerobic chemoheterotrophic conversions of PNSB along with the (an)aerobic chemoheterotrophic conversions of competing non-PNSB. These tests were conducted to understand how these conversions may individually contribute in a raceway reactor. The detailed cultivation conditions are described in Supporting Information S1

To explore the conversions of PNSB, four different conditions were tested under axenic conditions: (i) illumination with oxygen supply to study the combined photo- and chemotrophic growth (conditions prevalent in a raceway reactor); (ii) illumination without oxygen supply to study the phototrophic growth; (iii) no illumination with oxygen supply to study the aerobic chemotrophic growth and (iv) no illumination without oxygen supply to study the anaerobic chemotrophic growth (i.e. acidogenic metabolism).

An experiment was also performed to assess the effect of oxygen supply on the photo- and chemoheterotrophic growth of PNSB. The methodology is explained in Supporting Information S2.

#### 2.2.2 Light and oxygen availability as control tools to steer PNSB selectivity

These experiments were performed to explore the effects of light, oxygen supply, and the combination of light and oxygen on the biomass growth, biomass yield, biomass composition and PNSB selectivity.

A 100-L raceway reactor (MicroBio Engineering Inc., USA) was used to perform growth experiments in batch regime under non-axenic conditions. The stirring speed of the paddle was set at 30 rpm and the pH was controlled at 7 by sparging CO_2_. The maximum oxygen transfer rate (OTR) at this stirring speed was 204 mg O_2_ L^-1^ d^-1^. Temperature was controlled at 28°C with an external TetratecHT 300-W heater (Tetra, Germany). Approximately 5 L of tap water was added daily to compensate for evaporation. One halogen lamp was used to illuminate the raceway reactor at an intensity of 54 W m^-2^ (illumination spectrum see Supporting Information S4). The reactor was filled with the VFA-based medium and *Rhodobacter capsulatus* was used as inoculum at a total suspended solids (TSS) concertation of 0.02 g L^-1^.

Four different conditions were tested in batch: (i) 12-h light/12-h dark with 24-h stirring at a surface-to-volume ratio of 5 m^2^ m^-3^ as benchmark (reactor filled up to 100 L); (ii) 24-h light/0-h dark with 24-h stirring at a surface-to-volume ratio of 5 m^2^ m^-3^ to study the effect of light; (iii) 12-h light/12-h dark with 12-h stirring (reduced oxygen supply vs. 24-h stirring) during the light period at a surface-to-volume ratio of 5 m^2^ m^-3^ to study the effect oxygen supply and (iv) 12-h light/12-h dark with 24-h stirring at a surface-to-volume ratio of 10 m^2^ m^-3^ to study the effect of light (reactor filled up to 50 L). The absorbance of the biomass suspension (660 nm) was monitored to determine the maximum specific growth rate. Experiments were stopped when the stationary phase was reached. Samples were then taken for further analysis.

#### 2.2.3 Effect of SRT on PNSB growth

These experiments were performed to explore the effect of SRT on the productivity, biomass yield, biomass composition and PNSB selectivity.

Experiments were performed under combined photo- and chemotrophic conditions, allowing the entry of oxygen along with illumination (i.e. conditions prevalent in raceway reactor). Flasks of 500 mL were used as reactors and illuminated through a natural 12-h light/12-h dark regime with two halogen lamps at a light intensity of 30 W m^-2^ (vs. previous flask experiments section 2.2.1: 24-h light or 24-h dark). The flasks were filled with 200 mL of medium (section 2.1) corresponding to a maximum OTR of 336 mg O_2_ L^-1^ d^-1^. The experiment was performed non-axenically with *Rhodobacter capsulatus* as initial inoculum. The tested SRT were 1.25 d, 2 d and 3 d. All tests were performed in biological duplicate. After adding the medium and the inoculum, flasks were placed on a multipoint stirrer at 300 rpm.

Temperature during the light and dark period was respectively 29.3 ± 0.4°C (i.e. resulting from radiation heat) and 24.2 ± 0.7°C. The pH was 7.4 ± 0.2. Bottles were weighed daily to adjust for evaporation, remove a part of the broth and add fresh medium. Samples were taken daily to measure the absorbance (660 nm), pH, temperature and DO concentration. The moving average with a fixed subset size of three was determined. Steady state was reached when the daily absorbance overlapped with the moving average. Samples were then taken after the light and dark periods three days in a row to account for variability over time.

#### 2.2.4 Operational strategies to steer PNSB selectivity and reactor performance

A final experiment was performed to demonstrate that PNSB can be maintained in a raceway reactor over multiple generations and determine the best operational strategy in terms of productivity and COD removal.

Temperature, pH and stirring were controlled as described in section 2.2.2. Three operational strategies were tested: (i) 12-h light/12-h dark with 24-h stirring at a surface-to-volume ratio of 5 m^2^ m^-3^ as benchmark (reactor filled up to 100 L and depth 20 cm); (ii) 12-h light/12-h dark with 12-h stirring (reduces oxygen supply vs. 24-h stirring) during the light period at a surface-to-volume ratio of 5 m^2^ m^-3^ to study the effect oxygen supply and (iii) 12-h light/12-h dark with 24-h stirring at a surface-to-volume ratio of 10 m^2^ m^-3^ (higher light availability vs. 5 m^2^ m^-3^ reactor filled up to 50 L and depth 10 cm) to study the effect of light. The SRT was chosen based on the maximal specific growth rate during the batch experiments (Figure 2). A value of 0.8 d^-1^ was observed, which corresponds to a doubling time of 1.2 d. For safety reason, a SRT of 2 d was chosen for the three conditions to prevent washout from the reactor. Effluent was first removed and influent was then added before the start of the light period. The absorbance was analyzed daily. After steady state, samples were taken by the end of the light and dark periods three days in a row to account for variability over time. Samples were stored at −20°C for further analysis.

### 2.3 Analytical procedures

Standard methods were used to determine the TSS and volatile suspended solids concentration.^25^ The COD was measured using test kits (Macherey-Nagel, Germany) according to the manufacturer’s instructions. The volumetric mass transfer coefficient (KLa) of oxygen was determined according to the sulfite oxidation method.^26^ Protein concentration was analyzed by Markwell, et al. 1978^27^, which is adapted from the Lowry procedure. The bacteriochlorophyll a content was determined by an acetone/methanol solvent (7:2 v/v). extraction.^28^ Dissolved oxygen (DO) concentration (Hach, USA) and pH (Hanna Instruments, USA) were determined with Handheld meters.

### 2.4 Microbial community analyses

Genomic DNA was extracted from biomass samples collected (after steady state) across the reactor experiments using the DNeasy UltraClean microbial extraction kit (Qiagen, Venlo, the Netherlands) according to the manufacturer’s instructions. The DNA extracts were sent to a commercial company (Novogene, China) for amplicon sequencing analysis. In brief, the V3-V4 hypervariable region of the bacterial 16S rRNA gene pool of the DNA extracts was amplified by PCR using the pair of 341f/806r primers prior to sequencing of PCR products using a HiSeq 2500 sequencer (Illumina, USA). A detailed description of the wet-lab and dry-lab workflows can be found in Supporting Information S5.

### 2.5 Statistical analyses

Statistical analyses were performed in R (version 3.4.1) using RStudio (RStudio®, USA) for Windows.^29^ Student’s t-test were conducted to compare means. Normality of data residuals was tested using the Shapiro-Wilk normality test. The assumption of homoscedasticity was verified through a Levene’s test. The non-parametric Kruskal-Wallis rank sum test was executed when normality was rejected. The Welch’s t-test was used in case of heteroscedasticity. A significance level of *p* < 0.05 was chosen.

### 2.6 Production cost estimation

The cost of four PNSB production scenarios was estimated and compared: (i) a closed tubular PBR with 24-h stirring; (ii) a raceway reactor operated with 24-h stirring and a surface-to-volume ratio of 5 m^2^ m^-3^; (iii) a raceway reactor operated with 24-h stirring and a surface-to-volume ratio of 10 m^2^ m^-3^ and (iv) a raceway reactor operated with 24-h stirring during the light period and a surface-to-volume ratio of 5 m^2^ m^-3^. Illumination for the four scenarios was considered to be from sunlight according to a natural 12-h light/12-h dark regime For the experiments, synthetic wastewater was used. This cost estimation was performed with brewery wastewater, as a suitable model for food and beverage effluents where fecal contamination can be avoided.

The goal of this model was to compare a closed PBR with a raceway reactor operated at three different strategies. It was not intended to determine an accurate production cost for PNSB. This cost estimation ought to be seen as a decision-making tool for research. Details and all input parameters are presented in Supporting Information S6.

## 3 Results and discussion

### 3.1 Assess the conversions of PNSB and non-PNSB

The results of the individual conversions of PNSB (Figure 1) indicated that during the combined photo- and chemotrophic growth (i.e. light and oxygen supply), the phototrophic metabolisms (i.e. under light) was dominant and not their chemotrophic metabolism (i.e. oxygen supply). Biomass yields for their phototrophic and the combined photo- and chemotrophic metabolisms were similar (*p* > 0.05) and growth rates were almost equal. There was, therefore, more photo-assimilation of COD than oxidation to CO_2_. The bacteriochlorophyll content was lower for the combined photo- and chemotrophic metabolism than for the phototrophic metabolism, yet still 6 times higher than for the aerobic chemotrophic growth (i.e. no illumination with oxygen supply). It should be noted that the chemotrophic metabolism of PNSB contributes more to growth when the oxygen supply increases (Supporting Information S3).

**Figure 1.**
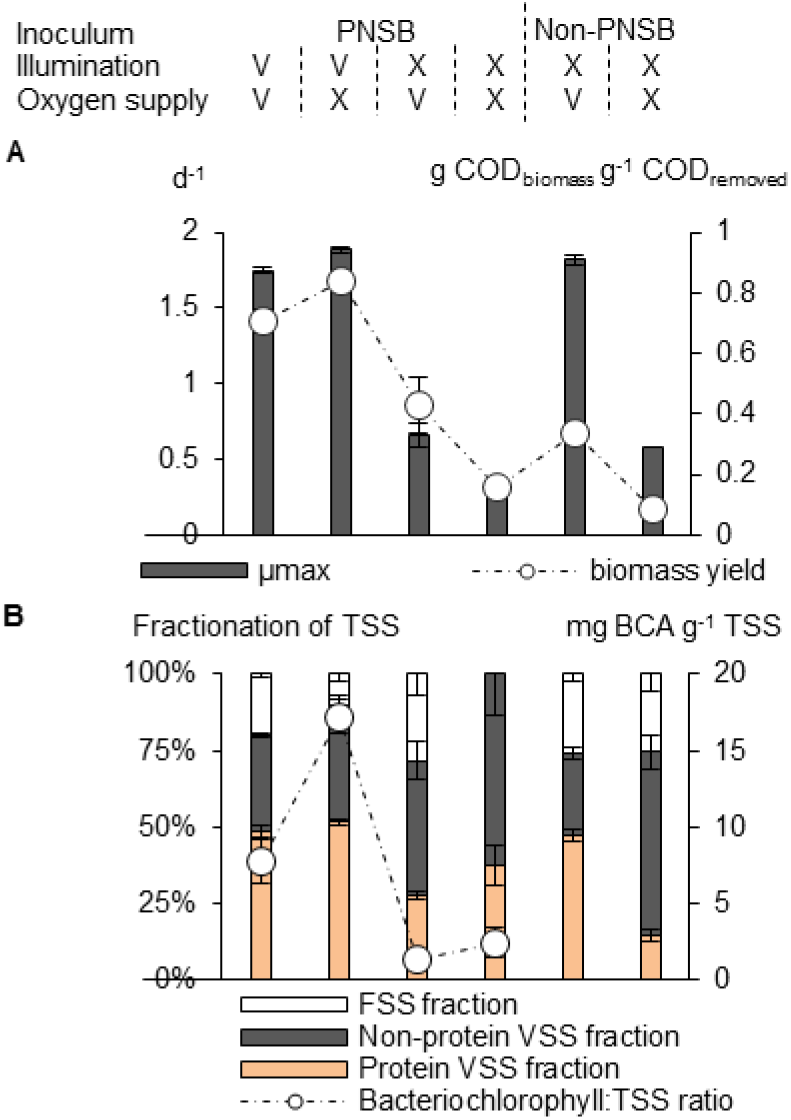
(A) maximum specific growth rate and biomass yield for purple non-sulfur bacteria (PNSB) and non-PNSB along with (B) biomass fractionation and bacteriochlorophyll a (BCA) content. Tested conditions: combined photo- and chemoheterotrophic (illumination: V; oxygen supply: V), photoheterotrophic (V; X), aerobic chemoheterotrophic (X; V) and anaerobic chemoheterotrophic (X; X) growth. The oxygen transfer rate was 336 mg O_2_ L^-1^ d^-1^. Experiments were performed axenically with *Rhodobacter capsulatus* used as model PNSB. Non-PNSB were grown non-axenically. Averages with standard error. TSS: total suspended solids; VSS: volatile suspended solids; FSS: fixed suspended solids i.e. ash

A similar experiment was performed by Sasaki, et al. 1998^30^ with *Rhodobacter sphaeroides* on orange peel waste, showing a 1.6 times higher biomass yield for the combined metabolism compared to the chemotrophic metabolism. Our results were in line with this finding. This reveals that PNSB have the potential to use both their photo- and chemotrophic metabolism at once.

The DO concentration in a raceway reactor is zero due to the direct consumption of oxygen, allowing anaerobic fermentation of COD. The anaerobic chemotrophic growth of PNSB was, therefore, tested with an organic more complex medium (vs. VFA-based medium see Supporting Information S1). PNSB were able to anaerobically ferment organics (Figure 1), yet growth rates and biomass yields were relatively low (0.3 ± 0.08 d^-1^; 0.16 ± 0.09 g COD_biomass_ g^-1^ COD_removed_). A similar observation was made by Schultz Weaver 1982^31^ with low anaerobic growth rates and biomass yields for *Rhodobacter capsulatus* (≈ 0.08 d^-1^; 0.09 g COD_biomass_ g^-1^ COD_removed_) and *Rhodospirillum rubrum* (≈ 0.13 d^-1^; 0.11 g COD_biomass_ g^-1^ COD_removed_). The non-PNSB inoculum showed growth rates of 0.58 ± 0.03 d^-1^ or 2 times higher compared to PNSB. Therefore, anaerobic fermentation will mainly be performed by competing non-PNSB. Stronger competition during the light and dark period might arise from aerobic chemotrophic non-PNSB since their growth rate was 2.8 times higher than for the aerobic chemotrophic growth of PNSB and equal to the combined photo- and chemotrophic growth (*p* > 0.05).

### 3.2 Light and oxygen availability as control tools to steer PNSB selectivity

This experiment was set up to demonstrate that PNSB can be produced in a raceway reactor under non-axenic conditions and to investigate the effect of light, oxygen supply and the combination of both on PNSB selectivity.

The four tested conditions in Figure 2 showed an increase of the bacteriochlorophyll and carotenoid peaks during the batch growth experiments. Microbial community analysis confirmed these findings with PNSB abundances between 8-31% (Supporting Information S7). Hence, this research pioneers in demonstrating PNSB production in a raceway reactor without strict anaerobic conditions, thereby setting a precedent for future research. Lu, et al. 2019^32^ have also claimed to produce PNSB in a PBR at DO concentrations between 0.2-0.5 mg O_2_ L^-1^, yet have not presented results of the microbial community composition.

**Figure 2.**
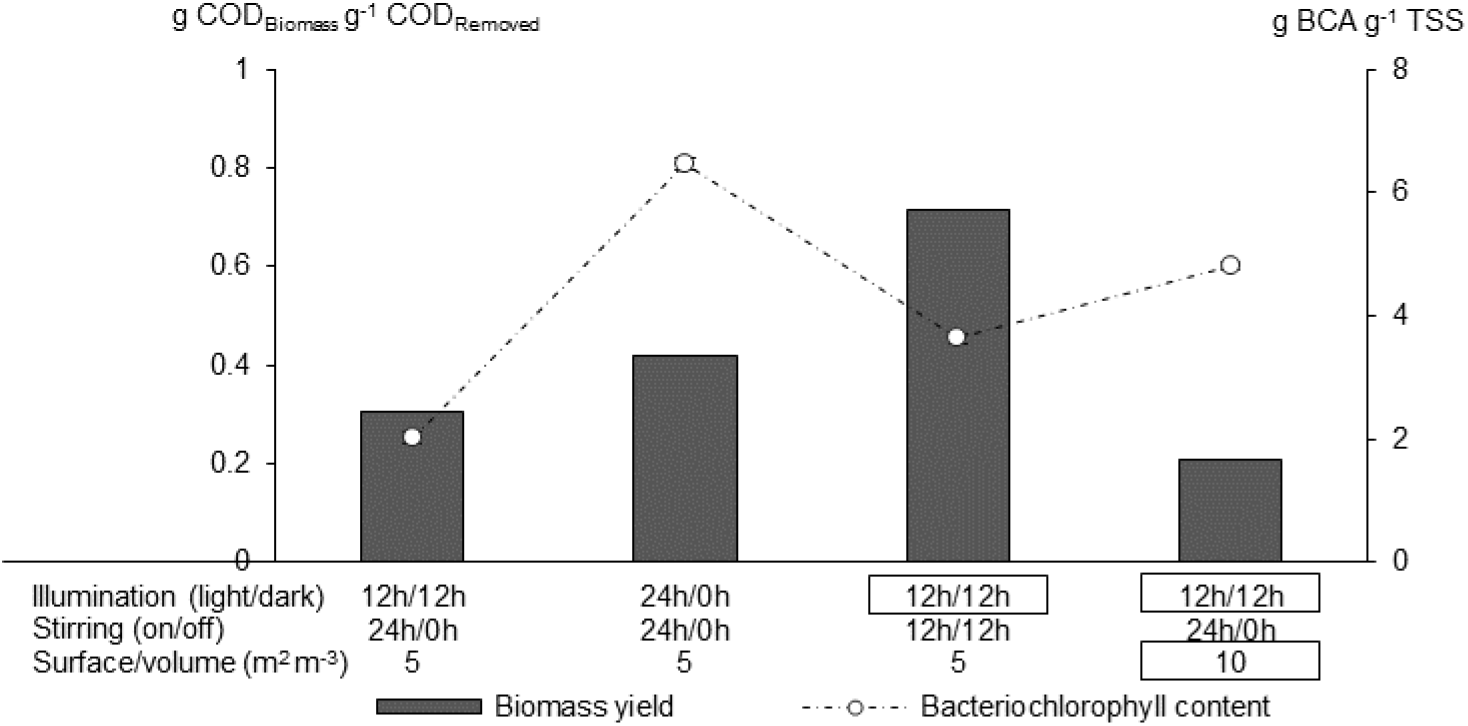
Batch raceway reactor experiment testing the effects of light (illumination), oxygen (stirring) and the combination of light (surface-to-volume ratio) on the biomass growth. Results show biomass yield (left y-axis) and bacteriochlorophyll a (BCA) content (right y-axis). Experiments were performed non-axenically with *Rhodobacter capsulatus* as initial inoculum. Stirring (on/off) 12 h / 12 h implies stirring during the light period and not during the dark. Rectangular boxes show the change in reactor operation relative to the control. TSS: total suspended solids

For light availability as control tool, increasing the illumination time to 24-h light/0-h dark (vs. benchmark 12-h light/12-h dark) resulted in a biomass yield increase of 1.4 times (more photo-assimilation), an increase of the bacteriochlorophyll content by 3.2 times and increase of the PNSB abundance by 3.9 times (31% vs. benchmark 8%; Supporting Information S7). This was also the most effective control tool in terms of PNSB abundance.

Preventing oxygen supply during the dark phase (not stirring vs. benchmark 24-h stirring) increased the biomass yield by 2.3 times, the bacteriochlorophyll content by 1.8 times and the PNSB abundance by 2.5 times (20%; Supporting Information S7). Hence, these results reconfirm the findings of Supporting Information S3, where a lower oxygen supply resulted in increased phototrophic growth. Increasing the surface-to-volume ratio had a dual effect. Relatively to the benchmark, the biomass yield decreased by 1.5 times due to increased COD oxidation and the bacteriochlorophyll content increased by 2.4 due to an increased light availability per reactor volume.

### 3.3 Effect of SRT on PNSB growth

This experiment was performed to study the effect of SRT on PNSB selectivity. Overall, PNSB abundances did not show substantial differences between SRT (Figure 3). This is in line with our previous observation where we tested the effect of SRT on PNSB abundance in a closed PBR.^13^ Notably (Figure 3), the PNSB abundance at 6 g COD L^-1^ for all SRT (50-67%) was lower compared to 3 g COD L^-1^ (88-94%). The exponent of the Shannon diversity index was also lower at 3 g COD L^-1^ (1.4-1.7) compared to 6 g COD L^-1^ (2.7-3.0), which indicates a more uneven community at lower loading rates. The abundance of the (an)aerobic chemoheterotrophs *Arcobacter* was predominantly higher at higher loading rates (17-36%). Overall, it can be concluded that a higher loading decreased the PNSB selectivity. This decreased PNSB selectivity was probably due to the higher COD availability especially during the night period, leading to increased growth of competing chemotrophs. As such, COD availability during the night will negatively influence the PNSB abundance.

**Figure 3.**
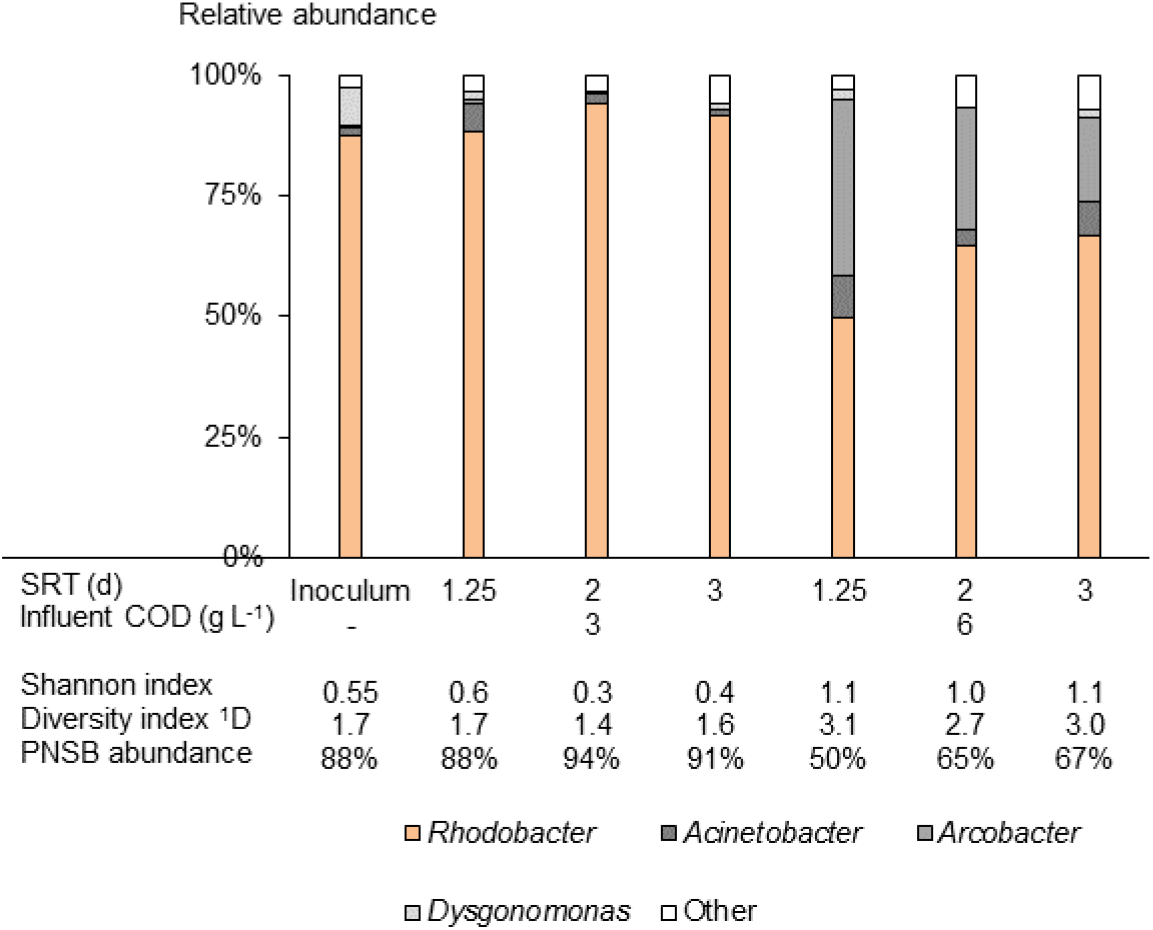
Effect of sludge retention time (SRT) on microbial community composition, Shannon’s H index, exp(H’) and purple non-sulfur bacteria (PNSB) abundance. Flasks were used as a reactor. The PNSB genera *Rhodobacter* and *Rhodopseudomonas* are all marked in orange. Samples obtained after 2-10 SRT.

### 3.4 Reactor performance and community dynamics over sequential batches

This experiment was conducted to demonstrate that PNSB can be maintained in a raceway reactor over multiple generations and determine the best operational strategy in terms of productivity and COD removal.

The highest productivities (0.21 g protein L^-1^ d^-1^ corresponding with 0.43 g TSS L^-1^ d^-1^) and removal rates (0.79 g COD L^-1^ d^-1^) were achieved when the reactor was operated with 24-h stirring and 12-h light/12-h dark at the highest surface-to-volume ratio of 10 m^2^ m^-3^ (Figure 4). A higher ratio of 10 m^2^ m^-3^ increased the light availability, resulting in higher biomass concentrations (0.81 ± 0.04 g TSS L^-1^) relative to the benchmark of 5 m^2^ m^-3^ (0.62 ± 0.02 g TSS L^-1^). For a closed PBR operated on the same medium at a SRT of 1 d (vs. 2 d for raceway reactor), we reached TSS productivities that were 1.5-2.6 times higher compared to the raceway reactor.^13^ This was probably due to the higher light availability in the PBR compared to a raceway reactor (surface-to-volume ratio 33 vs. 5-10 m^2^ m^-3^). For the microalga *Chlorella vulgaris* cultivated in the same reactor (12-h light/12-h dark), the productivity was 0.009 g protein L^-1^ d^-1^ or 22 times lower compared to PNSB.^6^ This was probably due to the higher growth rates of PNSB of 0.6-3.7 d^-1^ compared to the ones of microalgae of 0.60-1.38 d^-1^.^13,33,34^ To a lesser extent, non-PNSB chemotrophs also contributed to biomass production in the raceway reactor, thereby increasing the overall biomass productivity.

**Figure 4.**
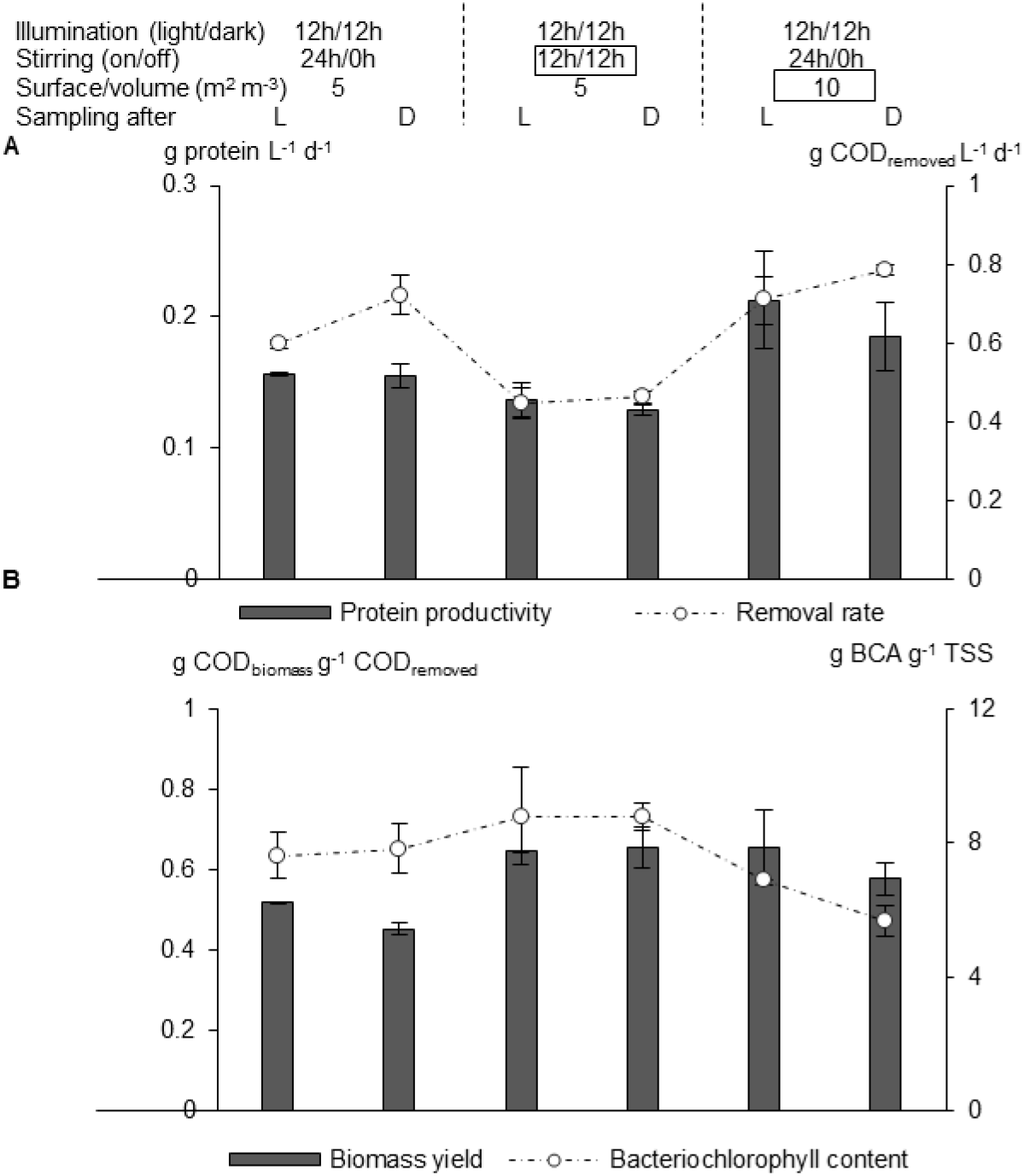
Production features of a raceway reactor operated at a sludge retention time of 2 d, testing the effect of oxygen (stirring) and light (surface-to-volume ratio). Sampling occurred after the light (L) or dark (D) period. Results show (A) the protein productivity and the volumetric removal rate along with (B) the biomass yield and bacteriochlorophyll (BCA) content. Experiments were performed non-axenically with *Rhodobacter capsulatus* as initial inoculum. Stirring (on/off) 12 h / 12 h implies stirring during the light period and not during the dark. Average values with standard error. 12 h / 0 h stirring occurred during the light period. Rectangular boxes show the change in reactor operation relative to the benchmark. TSS: total suspended solids

In terms of PNSB selectivity (Figure 5), preventing the combination of oxygen supply and darkness (not stirring) was an effective strategy in line with the results of section 3.2. The PNSB abundance was 56% (vs. 14% benchmark 24-h stirring) and the microbial community was more uneven showing a lower exponent of the Shannon diversity index (3.7 vs. 4.3 for benchmark). The decrease in PNSB abundance to 41% was notable during the dark period along with the increase of the exponent of the Shannon diversity index from 3.7 to 7.2. Hence, biomass harvesting should preferably be performed after the light period to assure a selective microbial community.

**Figure 5.**
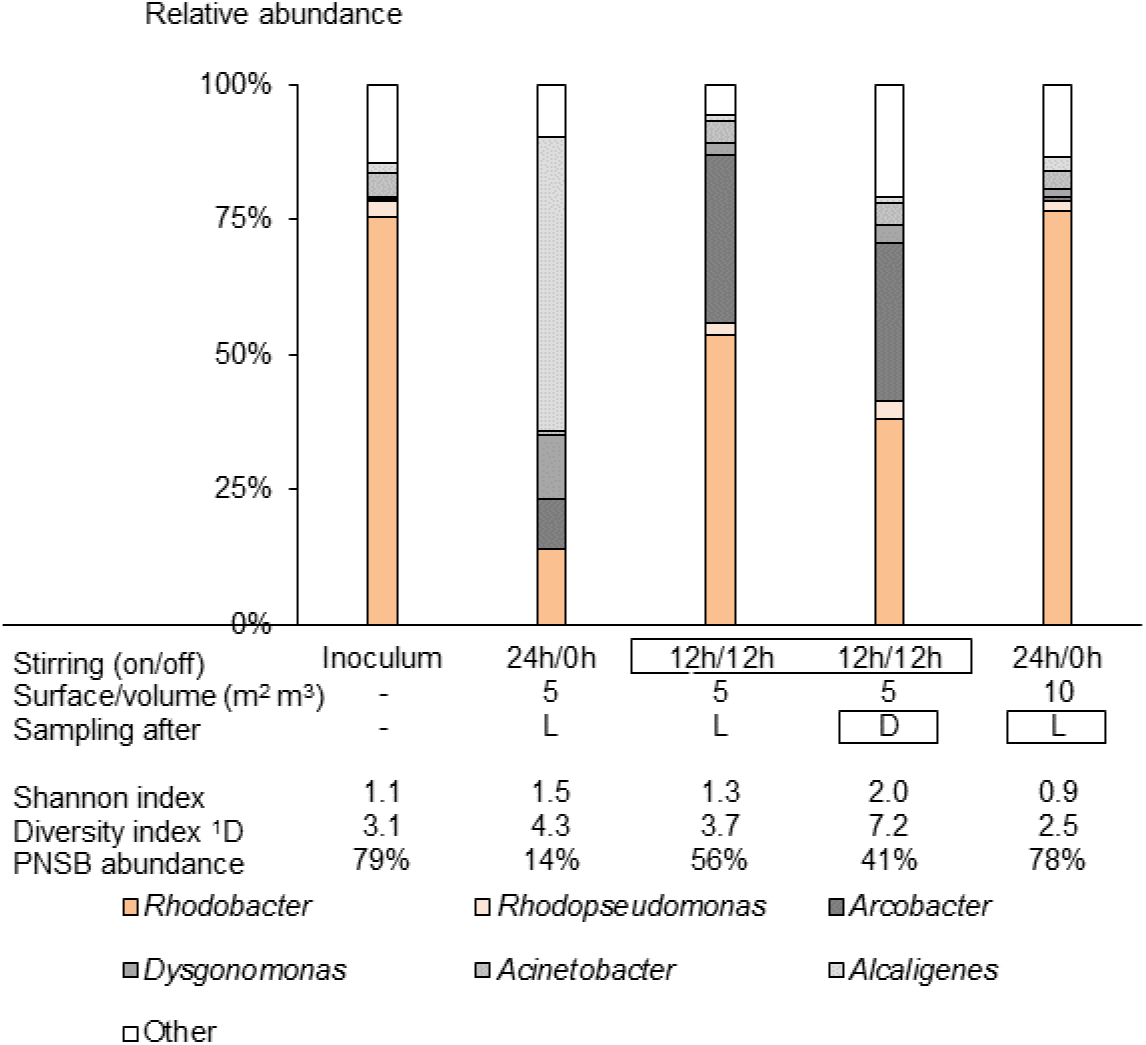
Effect of operational strategy of a raceway reactor on microbial community composition, Shannon’s H’ index, exp(H’) and purple non-sulfur bacteria (PNSB) abundance. Raceway reactor operated at a sludge retention time of 2 d, testing the effect of oxygen (stirring) and the combination of light (surface-to-volume ratio). Sampling occurred after the light (L) or dark (D) period. Stirring (on/off) 12 h / 12 h implies stirring during the light period and not during the dark. The PNSB genera *Rhodobacter* and *Rhodopseudomonas* are marked in orange. Rectangular boxes show the change in reactor operation relative to the benchmark.

The reactor operated with 24-h stirring at a surface-to-volume ratio of 10 m^2^ m^-3^ was the best strategy in terms of PNSB selectivity, showing a PNSB abundance of 78% or 5.6 times higher compared to the benchmark and very comparable to the inoculum. The exponent of the Shannon diversity index was only 2.5, the lowest for all conditions and even lower than the inoculum (3.5). This implies that light availability is key to boost PNSB selectivity in a raceway reactor. The findings also show that a raceway reactor can approach the PNSB selectivity of a closed PBR. Potential higher PNSB abundances might even be achieved if oxygen supply is prevented during the night along with high surface-to-volume ratios.

Other PNSB genera were also present in the system such as *Rhodopseudomonas* (2-3%), and *Blastochloris* (< 0.2%). The main competing genera were *Acinetobacter* (aerobic chemoheterotroph), *Dysgonomonas* (anaerobic chemoheterotrophs), *Arcobacter* ((an)aerobic chemoheterotrophs) and *Alcaligenes* (aerobic chemoheterotrophs) with an abundance of respectively 1-4%, 1-12%, 0-31% and 1-54%. Microalgae were not detected through the absorbance spectra (no chlorophyll peaks) and no cyanobacteria were identified by amplicon sequencing. This was probably due to the short SRT (2 d), resulting in washout of slower-growing microalgae (μ_max_ 0.60-1.38 d^-1^).^33^ Although the sulfate concentration in the medium was 1.6 g L^-1^, there were no SRB detected (of note, primers not designed for archaea). SRB require 0.7 g COD to remove 1 g of sulfate.^35^ Therefore, they can contribute to COD removal, yet biomass production will be negligible due to their low biomass yields of 0.015-0.033 g VSS g^-1^ SO4^2-^.^36^

According to the authors’ knowledge, this study is the first to publish results on PNSB production in a raceway reactor dedicated to microbial protein. Literature studying microbial communities in waste lagoons are helpful for benchmarking, as these systems enable photosynthesis and are open to air. Do, et al. 2003^37^ have investigated the correlation of environmental factors on photosynthetic blooms, i.e. the spontaneous growth of purple bacteria in waste lagoons. Their research showed that for a swine waste lagoon, up to 10% of the microbial community was made by a population of *Rhodobacter*. The authors observed a positive correlation between *Rhodobacter* and the sulfate concentration. They claimed that it was due to competition between PNSB and SRB. More PNSB growth would result in a lack of VFA for SRB and the accumulation of sulfate.

Our reactor was operated with artificial 12h-light/12h-dark conditions at constant temperature with a VFA-based medium. Hence, sunlight-driven pilot-scale validation should still confirm these findings. As scale-up for raceways is mainly horizontal (depth is kept constant), the main effects are expected from a different influent composition (more complex COD mixture vs. synthetic), fluctuations in COD flow and temperature, day/night length and light intensity. Another parameter that might be different for a for a full-scale raceway reactor is the oxygen supply to the system. The oxygen supply rate during the experiment in this chapter was around 204 mg O_2_ L^-1^ d^-1^. However, for a full-scale system this would potentially be lower due to higher reactor volume. The lower oxygen supply would positively influence the PNSB growth, yet might enhance the competition with anaerobic heterotrophs such as SRB and fermentative microorganisms.

### 3.5 Production cost comparison of a closed PBR and an open raceway reactor

An anaerobic reactor such as a closed PBR is the best non-axenic technology to selectively produce PNSB on wastewater, yet production costs were estimated between 2.7-6.4 times higher compared to a raceway reactor (Figure 6). These high costs can mainly be contributed to the high investment costs for a tubular PBR (47% of capital expenditure; CAPEX) and the energy required for recirculation (23% of operational expenditure; OPEX).

**Figure 6.**
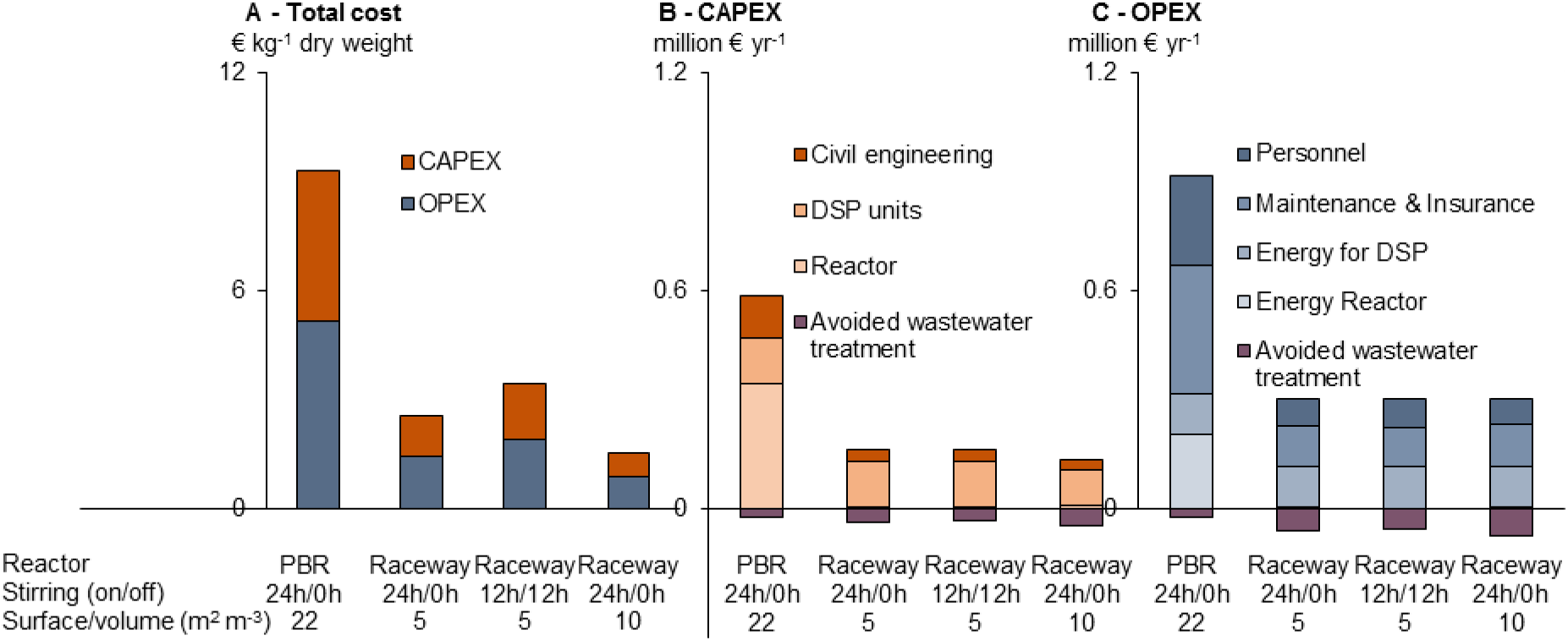
Cost comparison for a closed tubular photobioreactor (PBR) and an open raceway reactors using natural sunlight (12-h light/12-h dark). Results show (A) capital and operational expenditure (CAPEX, OPEX), (B) fractionation of CAPEX, and (C) fractionation of OPEX. The cost category “Avoided wastewater treatment” refers to activated sludge treatment costs that can be avoided due to COD removal by the microbial protein reactor. Stirring (on/off) 12 h / 12 h implies stirring during the light period and not during the dark. All scenarios were dimensioned based on brewery wastewater with a COD concentration of 2.1 kg m^-3^ and a flow rate of 1150 m^3^ d^-1^. DSP: Downstream processing i.e. harvesting, dewatering and drying.

PNSB cultivated in closed PBR have high biomass yields (1 g COD_biomass_ g^-1^ COD_removed_) due to photo-assimilation instead of oxidation of COD to CO_2_. This will result in lower COD removal efficiencies compared to a raceway reactor. A raceway reactor can save around €83,000-120,000 yr^-1^ of the wastewater treatment cost or 1.7-2.4 times more compared to a PBR. Raceway reactors have the potential to remove 88% of the incoming COD. Therefore, wastewater treatment after PNSB production with a raceway reactor can be a simple aerobic activated sludge treatment process instead of a digester followed by aerobic treatment. On the contrary, PBR only remove 35% of the COD. Hence, a digester is still needed to lower the COD concentration.

The cost to produce PNSB in a raceway reactor was estimated between €1.3-3.0 kg^-1^ dry weight (DW). Downstream processing was the most dominant cost factor contribution by 57-61% to the CAPEX and by 40-43% to the OPEX. A further cost reduction of 27-40% can be possible if ultrafiltration is replaced by gravitational settling (€0.9-2.4 kg^-1^ DW). Currently, there is no published literature studying the settleability of PNSB except for Cerruti, et al. 2020 DOI: https://doi.org/10.1101/2020.01.08.899062^38^. Further research should explore this closely, as it will be an important cost saver.

The operational strategy with 24-h stirring at a surface-to-volume ratio of 10 m^2^ m^-3^ (reactor depth 10 cm) resulted in the highest PNSB selectivity and was also the most cost-effective. A regime of 24-h stirring will produce more biomass (more chemotrophic growth) and will remove more COD relatively to 12-h stirring. This will result in a cost that can be spread out over more products and additional savings for the subsequent aerobic activated sludge process.

## 4 Conclusions

The goal of this research was to develop control tools to selectively produce PNSB in a raceway reactor. The main findings of this study are:

i. This study pioneers in the selective production of PNSB in a raceway reactor with productivities of up to 0.43 g TSS L^-1^ d^-1^ and COD removal rates of up to 0.79 g COD L^-1^ d^-1^.
ii. Avoiding oxygen supply during the dark phase and a higher surface-to-volume ratio were the best operational strategies to maximize the PNSB abundance (56-78%) and lower the diversity.
iii. SRT does not show to have an impact on PNSB selectivity. However, COD availability should be avoided in the dark, as it decreases the PNSB abundance from 90% to 69% and increases the Shannon diversity from 0.45 to 1.1.
iv. Flask and raceway experiments showed that PNSB competed mainly with aerobic chemotrophs and to a minor extent with anaerobic chemotrophs. Microalgae or SRB were not identified as major competitors.
v. The combination of oxygen supply, higher COD load and darkness should be avoided.
vi. Production costs for a raceway reactor amount to €1.9 kg^-1^ DW, which is six times cheaper than a closed PBR.

## Acknowledgments

The authors kindly acknowledge (i) the Research Foundation Flanders (FWO-Vlaanderen) for supporting A.A. with a doctoral fellowship (strategic basic research; 1S23018N), (ii) the Rosa Blanckaert Foundation for supporting A.A with a research grant, (iii) the Belgian Science Policy Office for their support to MELiSSA (CCN5 to C4000109802/13/NL/CP), (iv) ESA’s life support system R&D program, which scientifically and logistically supported this study (http://www.esa.int/Our_Activities/Space_Engineering_Technology/Melissa), and (v) Matthijs Juchem and Enerelt Bilegt for their assistance with the raceway reactor experiments. D.G.W. and M.C. are supported by a start-up grant of the Department of Biotechnology of the TU Delft.

