## Supporting Information for "Control tools to selectively produce purple bacteria for microbial protein in raceway reactors"

#### S1. Cultivation conditions

*Rhodobacter capsulatus* was first axenically pre-cultivated in a climate chamber (Snijders Scientific) with a pre-autoclaved VFA medium based on Alloul, et al. 2019[1]. Flasks of 500 mL were then filled with 200 mL of autoclaved VFA medium and inoculated with *Rhodobacter. capsulatus* at an initial total suspended solids (TSS) concentration of 0.02 g $L^{-1}$. The flasks had a surface-to-volume ratio of 61 $m^2$ $m^{-3}$ (illuminated surface divided by volume flask). PNSB are not able to anaerobically convert VFA in the dark.[2] Therefore, condition 'iv' was performed with a medium more complex in organic composition (Table S1). Flasks that were tested without illumination, were covered with an aluminum foil to prevent light penetration (conditions 'iii' & 'iv'). Flasks without oxygen supply were closed with a screw cap and flushed with nitrogen gas for 1 min (conditions 'ii' & 'iv'). All flasks were then placed on a multipoint stirrer at 300 rpm (Thermo Scientific, USA) in a climate chamber (Snijders Scientific) at 28°C equipped with fluorescent lamps (Sylvania, Germany) of 36 W at a light intensity of 30 W $m^{-2}$. All experiments were tested in triplicate. For the non-PNSB inoculum, only conditions 'iii' no illumination with oxygen supply and condition 'iv' no illumination without oxygen supply were tested. The maximum oxygen transfer rate (OTR) of the flasks with oxygen supply was 336 mg $O_2$ $L^{-1}$ $d^{-1}$ (condition 'i' and 'iii'). Growth was monitored by measuring the absorbance at 660 nm along with the bacteriochlorophyll absorbance peaks (800 nm and 860 nm) to confirm the presence of PNSB. Samples were taken at the start and the end of the experiment for further analysis.

**Table S1** Medium composition to study anaerobic chemotrophic growth of PNSB and non-PNSB

| Ingredient | Mass (mg $L^{-1}$) |
|---|---|
| Beer* | 94 |
| Yeast | 548 |
| Malt extract | 1565 |

| | |
|---|---|
| Peptone | 671 |
| $(NH_4)_2SO_4$ | 1570 |
| $MgSO_4$ | 90 |
| $CaCl_2$ | 75 |
| $FeSO_4.7H_2O$ | 34 |
| Acetic acid | 175 |
| Propionic acid | 154 |
| Butyric acid | 26 |

*Cara Pils beer (Colruyt, Belgium)

#### S2. Methodology: Influence of oxygen supply on PNSB growth

In order to use oxygen supply as control tool to steer the PNSB selectivity, it should first be understood how their individual photo- and chemotrophic growth are affected by oxygen. That is why the following experiments were conducted to assess the effect of oxygen supply on the photo- and chemoheterotrophic growth of PNSB.

Two metabolic conditions were tested at two oxygen transfer rates (OTR) levels of 72 and 336 mg $O_2$ $L^{-1}$ $d^{-1}$: (i) illumination with oxygen supply to examine the combined photo- and chemotrophic growth of PNSB and (ii) no illumination with oxygen supply to study the aerobic chemotrophic growth of PNSB. The experiment was performed axenically with the medium described in section 2.1 of the manuscript. Flasks of 500 mL were either filled with 200 or 500 mL of medium to achieve a maximum OTR of 336 or 72 mg $O_2$ $L^{-1}$ $d^{-1}$, respectively (OTR = -0.88 x volume + 512). The OTR of 336 mg $O_2$ $L^{-1}$ $d^{-1}$ was chosen, as it was in the same order of magnitude as the OTR of a raceway reactor (204 mg $O_2$ $L^{-1}$ $d^{-1}$). All flasks had a surface-to-volume ratio of 61 $m^2$ $m^{-3}$. Flasks that were tested without illumination, were covered with an aluminum foil to prevent light penetration. Triplicate flasks were then placed on a stirrer and monitored as described in S1.

#### S3. Influence of oxygen supply on PNSB growth

Increasing the oxygen supply for the combined photo- and chemotrophic growth of PNSB, shifted the dominance from phototrophic to chemotrophic growth (Figure S1). This was evident from the following observations: (i) growth rates for the combined photo- and chemotrophic metabolism were equal ($p > 0.05$), yet the aerobic chemotrophic growth of PNSB increased by 1.9 times. This implies that chemotrophic growth contributed more to the combined photo- and chemotrophic metabolism; (ii) the biomass yield for the combined photo- and chemotrophic was 1.4 times lower at the higher OTR and thus there was more COD oxidation to $CO_2$ instead of photo-assimilation; (iii) the bacteriochlorophyll content decreased by 1.7 times shifting more to the pigment level of chemotrophic growth.

These findings show that controlling the oxygen availability in a raceway reactor is crucial. A higher oxygen supply will not only increase the chemotrophic growth rate of PNSB, but also of competing chemotrophs that grow even faster than PNSB.

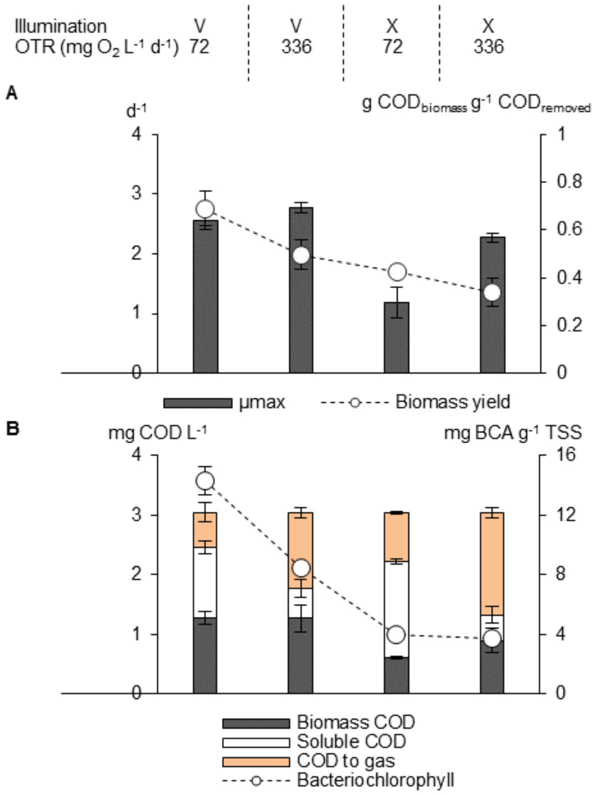

**Figure S1** Growth performance for purple non-sulfur bacteria (PNSB) grown under combined photo- and chemoheterotrophic (illumination: V) and aerobic chemoheterotrophic (X) conditions at two maximum oxygen transfer rates (OTR): (A) maximum specific growth rate and biomass yield along with (B) COD balance and bacteriochlorophyll a (BCA) content. COD to gas determined by subtracting total COD at the start of the experiment with biomass COD and soluble COD at the end of the experiment. Experiments were performed axenically with *Rhodobacter capsulatus* used as model PNSB. Averages with standard error.

#### S4. Illumination spectrum raceway reactor

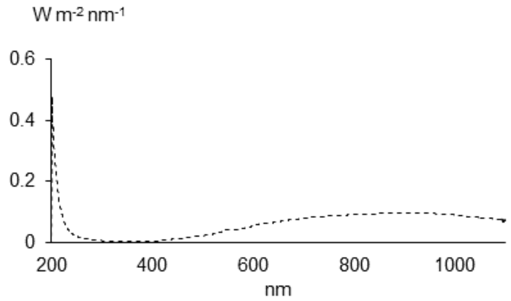

**Figure S2** Illumination spectrum of the halogen lamp used to illuminate the raceway reactor.

#### S5. Microbial community analyses

Genomic DNA was extracted using the DNeasy UltraClean microbial extraction kit (Qiagen, Venlo, the Netherlands) according to the manufacturer's instructions. The extracted DNA was sent to a commercial company (Novogene, Hong Kong, China) for amplicon sequencing analysis. The set of forward 341f (5'- CCTACGGGAGGCAGCAG-3') and reverse 806r (5'-GGACTACHVGGGTWTCTAAT-3') primers [3] were used to amplify the V3-V4 hypervariable region of the 16S rRNA gene by polymerase chain reaction (PCR). The amplicon sequencing libraries were pooled and sequenced in an Illumina paired-end platform. After sequencing, the raw reads were quality filtered, chimeric sequences were removed and OTU were generated on the base of ≥ 97% identity. Subsequently, microbial community analysis was performed by Novogene using Mothur & Qiime software (V1.7.0). For phylogenetical determination SSURef database from SILVA (http://www.arb-silva.de/) was used. Relative abundances of OTU were reported as % total sequencing reads count.

#### S6. Production cost estimation

The economic evaluation was based on a four-step methodology: (i) constructing a process flow diagram for the production and downstream processing of purple non-sulfur bacteria (PNSB), (ii) collecting literature based data (Table S1) for capital expenditure (CAPEX) and operational expenditure (OPEX) per process unit, (iii) creating the mass, water and energy balance for the whole process and (iv) calculating the CAPEX and OPEX for all four scenarios. Input parameters in Table S3 were used to calculated brewery treatment cost. The difference between wastewater treatment without and with PNSB production was subtracted from the PNSB production cost.

The volumetric flow rate, COD, total nitrogen, and total phosphorus concentration were respectively 1150 $m^3\ d^{-1}$, 2.11 kg $m^{-3}$, 0.04 kg $m^{-3}$ and 0.004 kg $m^{-3}$.[1] Nitrogen and phosphorus dosage was included, when more nutrients were converted to microbial biomass than present in the wastewater. The COD removal efficiency and COD removal rates for the three raceway reactor scenarios were based on the experiments of manuscript section 3.6. The SRT (2 d) was considered to be equal to the hydraulic retention time (no biomass recirculation). All scenarios were a series of subsequent processes: PNSB production in a tubular PBR or in a raceway reactor followed by ultrafiltration, centrifugation, and drying. Other parameters such as construction, piping, and mixing were also taken into account. PNSB removed between 32-88% COD and the avoided costs for the subsequent aerobic activated sludge treatment were also included in the cost model.

**in Table S2** Technological and economic input parameters for four production cost simulations: (I nu) a photobioreactor operated with 24h-stirring and a surface to volume ratio of 22 $m^2$ $m^{-3}$, (ii) a raceway reactor operated with 24h-stirring and a surface to volume ratio of 5 $m^2$ $m^{-3}$, (iii) a raceway reactor operated with 24h-stirring and a surface to volume ratio of 10 $m^2$ $m^{-3}$ and (iii) a raceway operated with12h-stirring during the light period and a surface to volume ratio of 5 $m^2$ $m^{-3}$. CAPEX: capital expenditure, OPEX: operational expenditure, COD: chemical oxygen demand, VSS: volatile suspended solids, TSS: total suspended solids, el: electrical energy, th: thermal energy, UF: ultrafiltration. * kg water removed

| | **CAPEX** | | | | **OPEX** | | | |
|---|---|---|---|---|---|---|---|---|
| **Process Unit** | **Parameter** | **Value** | **Unit** | **Reference** | **Parameter** | **Value** | **Unit** | **Reference** |
| General | Depreciation time | 20 | yr | 4 | Electricity cost | 0.11 | € $kWh_{el}^{-1}$ | 5 |
| | Interest rate | 1.67 | % | | Natural gas Europe | 0.03 | € $kWh_{th}^{-1}$ | 6 4 |
| | Construction/installation | 25 | % CAPEX | 7 | Working days | 260 | d $yr^{-1}$ | |
| | Piping/electrical | 30 | % CAPEX | | US Dollar to Euro | 0.84 | € $\$^{-1}$ | 8 |
| | Land | 10 | € $m^{-2}$ | 9 | Ammonium | 0.51 | € $kg^{-1}$ N | 10 |
| | | | | | Phosphate | 2.79 | € $kg^{-1}$ P | 9 |
| | | | | | Sludge yield PBR | 0.70 | g VSS $g^{-1}$ COD | 1 |

| | CAPEX | | | | OPEX | | | |
|---|---|---|---|---|---|---|---|---|
| **Process Unit** | **Parameter** | **Value** | **Unit** | **Reference** | **Parameter** | **Value** | **Unit** | **Reference** |
| | | | | | Sludge yield raceway | 0.36-0.41 | g VSS $g^{-1}$ COD | Experiments |
| | | | | | Operation | 1 | % CAPEX | [11] |
| | | | | | Maintenance | 1.5 | % CAPEX | |
| | | | | | Insurance | 0.3 | % CAPEX | |
| PBR | Cost PBR | 5000 | € $m^{-3}$ | [7] | Mixing energy | 2500 | W $m^{-3}$ | [4] |
| | Surface to volume ratio | 22 | $m^2$ $m^{-3}$ | [9] | | | | |
| | Circulation pump capacity | 1000 | $m^3$ $h^{-3}$ | | | | | |
| | Retention time circulation | 0.05 | h | | | | | |
| | Circulation pump | 2900 | € $unit^{-1}$ | | | | | |
| Raceway reactor | Liner | 4.9 | € $m^{-2}$ | [12] | Mixing energy | 3.72 | W $m^{-3}$ | |
| | Surface to volume ratio | 5 or 10 | $m^2$ $m^{-3}$ | [4] | | | | |
| | Paddlewheel | 22.45 | € $m^{-3}$ | [12] | | | | |

| | **CAPEX** | | | | **OPEX** | | | |
|---|---|---|---|---|---|---|---|---|
| **Process Unit** | **Parameter** | **Value** | **Unit** | **Reference** | **Parameter** | **Value** | **Unit** | **Reference** |
| | Landscaping | 0.94 | € m$^{-3}$ | [13] | | | | |
| UF | Cost UF | 178 | € unit$^{-1}$ | [12] | Energy membrane | 1.78 | kWh$_{el}$ m$^{-3}$ | [12] |
| | Capacity | 1 | m$^{3}$ d$^{-1}$ | | Biomass recovery | 100 | % | [4] |
| Centrifuge | Cost centrifuge | 6720 | € unit$^{-1}$ | [14] | Energy centrifuge | 1.35 | kWh$_{el}$ m$^{-3}$ | [12] |
| | Capacity | 4.8 | m$^{3}$ d$^{-1}$ | | Biomass recovery | 97 | % | [4] |
| | | | | | Heat demand | 1.75 | kWht$_{th}$ kg$^{-1*}$ | [15] |
| Dryer | Cost spray dryer | 460000 | € unit$^{-1}$ | [16] | Biomass recovery | 95 | % | [4] |
| | Capacity | 1.2 | m$^{3}$ d$^{-1}$ | | | | | |

**Table S3** Technological and economic input parameters for cost calculation of brewery wastewater treatment. CAPEX: capital expenditure, OPEX: operational expenditure, COD: chemical oxygen demand, VSS: volatile suspended solids, TSS: total suspended solids, el: electrical energy based on van Haandel van der Lubbe 2012[11]

| **CAPEX** | | | | **OPEX** | | | |
|---|---|---|---|---|---|---|---|
| **Parameter** | **Value** | **Unit** | **Reference** | **Parameter** | **Value** | **Unit** | **Reference** |
| Depreciation time structures | 20 | yr | [11] | Electricity cost | 0.11 | € $kWh_{el}^{-1}$ | [11] |
| Interest rate | 1.67 | % | [4] | Working days | 260 | d $yr^{-1}$ | [4] |
| Cost acidogenic tank | 189 | € $m^{-3}$ | [11] | US dollar to Euro factor | 0.84 | € $\$^{-1}$ | [8] |
| Cost UASB | 336 | € $m^{-3}$ | | Sludge concentration UASB | 17.5 | g TSS $L^{-1}$ | [11]) |
| Cost aerobic reactor | 189 | € $m^{-3}$ | | Anaerobic sludge yield | 0.05 | g VSS $g^{-1}$ COD | |
| Cost settler | 252 | € $m^{-3}$ | | Sludge age | 37 | d | |
| Cost thickener | 504 | € $m^{-3}$ | | Temperature | 30 | °C | |
| Cost aeration | 420 | € $kW^{-1}$ | | Ratio COD:VSS | 1.42 | g COD $g^{-1}$ VSS | |
| Additional construction cost | 35 | % CAPEX | | Organic sludge fraction (UASB) | 0.65 | g VSS $g^{-1}$ TSS | |

| **CAPEX** | | | | **OPEX** | | | |
|---|---|---|---|---|---|---|---|
| **Parameter** | **Value** | **Unit** | **Reference** | **Parameter** | **Value** | **Unit** | **Reference** |
| Additional investment costs | 1.6 | % CAPEX | | Aerobic sludge yield | 0.45 | g VSS $g^{-1}$ COD | |
| | | | | Organic sludge fraction aerobic reactor | 0.85 | g VSS $g^{-1}$ TSS | |
| | | | | Sludge concentration aerobic reactor | 5 | g VSS $L^{-1}$ | (Personal communication, Palm Breweries) |
| | | | | Sludge age | 20 | d | |
| | | | | Sludge disposal | 200 | € $tonne^{-1}$ TSS | [11] |
| | | | | Operation | 1 | % CAPEX | |
| | | | | Maintenance | 1.5 | % CAPEX | |

| CAPEX | | | | OPEX | | | |
|---|---|---|---|---|---|---|---|
| **Parameter** | **Value** | **Unit** | **Reference** | **Parameter** | **Value** | **Unit** | **Reference** |
| | | | | Insurance | 0.3 | % CAPEX | |
| | | | | Personnel | 2 | % CAPEX | |

### S7. Effect of control tools on the microbial community composition

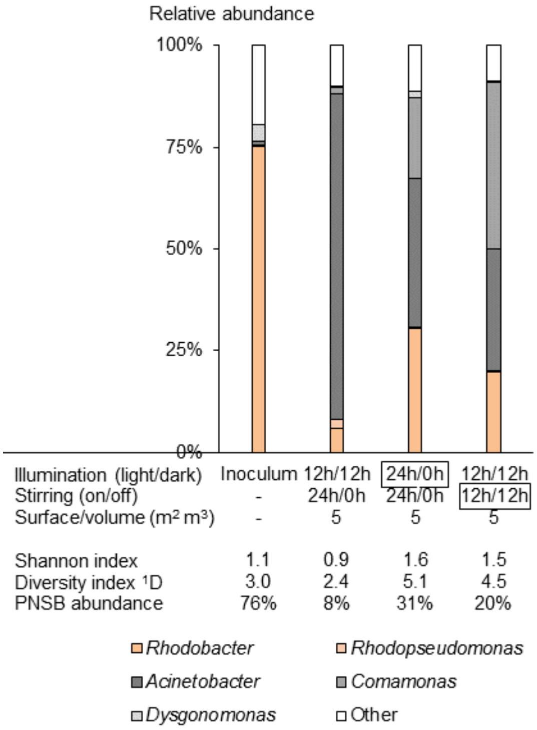

**Figure 3** Effects of light (illumination) and oxygen (stirring) on the microbial community composition, Shannon index, diversity index which is the exponential of the Shannon index and purple non-sulfur bacteria (PNSB) abundance. Experiments performed in a raceway reactor operated in batch regime. The PNSB genera *Rhodobacter* and *Rhodopseudomonas* are all marked in orange. Grey markings show the change in reactor operation relatively to the benchmark.

(5) Eurostat. Electricity prices for non-household consumers. http://appsso.eurostat.ec.europa.eu/nui/submitViewTableAction.do

(6) Eurostat. Gas prices for non-household consumers. http://appsso.eurostat.ec.europa.eu/nui/submitViewTableAction.do (accessed Dec 13, 2017).

(7) Acien, F. G.; Fernandez, J. M.; Magan, J. J.; Molina, E. Production cost of a real microalgae production plant and strategies to reduce it. *Biotechnol Adv* **2012,** *30*, 1344-1353.

(8) Google Inc. Google currency converter. www.google.be (accessed December 12, 2017).

(9) Norsker, N. H.; Barbosa, M. J.; Vermue, M. H.; Wijffels, R. H. Microalgal production - A close look at the economics. *Biotechnol Adv* **2011,** *29*, 24-27.

(10) Amer, L.; Adhikari, B.; Pellegrino, J. Technoeconomic analysis of five microalgae-to-biofuels processes of varying complexity. *Bioresour Technol* **2011,** *102*, 9350-9.

(11) van Haandel, A. C.; van der Lubbe, J. G. M. *Handbook of biological wastewater treatment: design and optimisation of activated sludge systems;* IWA Publishing: 2012.

(12) *Process design and economics for the production of algal biomass: algal biomass production in open pond systems and processing through dewatering for downstream conversion;* National Renewable Energy Laboratory: Denver, CO, 2016 https://www.nrel.gov/docs/fy16osti/64772.pdf.

(13) Rogers, J. N.; Rosenberg, J. N.; Guzman, B. J.; Oh, V. H.; Mimbela, L. E.; Ghassemi, A.; Betenbaugh, M. J.; Oyler, G. A.; Donohue, M. D. A critical analysis of paddlewheel-driven raceway ponds for algal biofuel production at commercial scales. *Algal Res* **2014,** *4*, 76-88.

(14) Li, J.; Zhu, D. L.; Niu, J. F.; Shen, S. D.; Wang, G. C. An economic assessment of astaxanthin production by large scale cultivation of *Haematococcus pluvialis*. *Biotechnol Adv* **2011,** *29*, 568-574.

(15) Baker, C. G. J.; McKenzie, K. A. Energy consumption of industrial spray dryers. *Dry Technol* **2005,** *23*, 365-386.

(16) Ólafsson, S. F. Downstream process design for microalgae. Master thesis, University of Iceland, 2013.
